# HA Nanocarriers Mediate HepG2 Vaccine Induction of hGM-CSF Gene Antitumor effect study

**DOI:** 10.1101/636316

**Authors:** Leah Robert, David Gupta

**Author notes:** Address: 250 Hackberry Ln, Tuscaloosa, AL 35401.

## Abstract

This report developed a novel method to observe the anti-tumor effect of HA nanoparticle carrier-mediated HepG2 cell vaccine transfected with hGM-CSF gene in vitro, and provide evidence for the clinical application of hGM-CSF gene-modified HepG2 cell vaccine. HA nanoparticle-mediated hGM-CSF gene transfection of HepG2 cells was used to prepare HepG2 cell vaccine transfected with GMCSF gene. Human PBMC were isolated by density gradient centrifugation and human PBMC were induced in vitro. The proliferative activity of PBMC and the killing effect on HepG2 cells were determined by WST-1 method. The positive expression rates of CD4+ and CD8+ were analyzed by flow cytometry, and the secretion of INF-γ was determined by ELISA. WST-1 results showed that the transgenic HepG2 vaccine induced PBMC proliferation, and its proliferation rate was better than that of wild-type vaccine. The induced PBMC had a higher killing rate against HepG2 than the wild-type vaccine group and each blank. In the control group, FCM results showed that the positive expression rates of CD4+ and CD8+ in the transgenic HepG2 vaccine group were higher than those in the wild-type vaccine group and each blank control group. The ELISA results showed that the IFN-γ content in the transgenic PBMC culture supernatant was 1989.76. +/− 254.21 pg/ml, higher than the wild-type vaccine group and each blank control group.

## Introduction

Primary liver cancer (PLC) is one of the common and refractory malignant tumors in China, 90% of which are hepatocellular carcinoma (HCC), accounting for the second highest mortality rate of malignant tumors in China. Surgical resection is the preferred method for the treatment of liver cancer, but its efficacy is still not satisfactory. The recurrence rate is as high as 60%-70% or even higher after 5 years [1]. Recurrence and metastasis of tumor remains a difficult problem to be solved. With the in-depth study of tumor immunity, tumor vaccines prepared by transferring cytokine genes with extensive regulatory functions to tumor cells have achieved encouraging results in many clinical and experimental studies [2] -4]. In this experiment, HepG2 cells were transfected with GM-CSF gene in HA nanocarriers and inactivated by 60Co intermittent irradiation to prepare HepG2 cell vaccine with hGM-CSF gene. Peripheral blood mononuclear cell (PBMC) was isolated, and the transgenic vaccine was co-cultured with PBMC to induce PBMC in vitro, and the proliferation, differentiation and killing effects of PBMC after induction were observed to evaluate its effect on immune function. To provide a basis for the clinical application of HepG2 cell vaccine transfected with hGMCSF gene.

## Materials and method

HepG2 cells were purchased from the Shanghai Institute of Cell Biology, Chinese Academy of Sciences. Transfer of hGM-CSF gene HepG2 cells were prepared from previous experiments and frozen in the South China University Cancer Institute. Fresh human blood is provided by healthy volunteers, aged 24 to 35 years old, with an average age of 28 years, 3 males and 2 females. The total RNA extraction kit, RT kit, and PCR kit were purchased from Sigma. Both the GAPDH and hGMCSF gene PCR primers were synthesized by Shanghai Biotech Co., Ltd. L glutamine was purchased from Beijing Dingguo Biotechnology Co., Ltd., recombinant GMCSF was purchased from PeproTech, USA, and recombinant human IL-2 was purchased from Jiangsu Jinsili Pharmaceutical Co., Ltd.; human peripheral blood lymphocyte separation solution (specific gravity 1.077)) purchased from Tianjin Yuyang Biological Co., Ltd.; anti-human FITC-CD4 mAb, anti-human PE-CD8 mAb, IgG2-FITC/IgG1-PE purchased from Ebioscience; human GMCSF ELISA test kit, human INF-γ ELISA test kit The interferon kit was purchased from Wuhan Boster Bioengineering Co., Ltd., and the WST-1 cytotoxicity and proliferation assay kit was purchased from Biyuntian Company.

The HA nanoparticle carrier mediates HepG2 cells transfected with hGM-CSF gene [5], and then combined with wild-type HepG2 cells by sublethal dose of radiation to prepare HepG2 vaccine and wild-type HepG2 vaccine transfected with GMCSF gene. . RT-PCR was used to identify the expression and integration of hGMCSF-mRNA after irradiation, and the secretion and duration of hGMCSF in culture medium were determined by ELISA.

Peripheral venous blood of healthy volunteers was taken under aseptic conditions, and human PBMC was isolated by density gradient centrifugation. Cell suspension was prepared by adding 1640 complete medium containing 10% FBS, and the cell density was adjusted to 1 × 10E6/ml. Place in a CO2 cell culture incubator (37 ° C, 5% CO 2, 95% humidity) for use.

The irradiated transgenic HepG2 vaccine and the irradiated wild-type HepG2 vaccine were adjusted with 1640 medium containing 15% FBS to adjust the cell concentration in 1640 medium to 2 × 10E5 / ml, 1 × 10E5 / ml, 5 × 10E4 / Ml, 2.5 × 10E4/ml. Add 1 × 10E6/ml PBMC to a 96-well plate at 100 μl per well. 100 μl of each of the abovementioned stimulator cells was added to each well, and the ratio of the stimulating cells to the PBMC cells was 1:40, 1:20, 1:10, 1:5, respectively, and 5 wells for each ratio, all culture wells. Add 0.5 μl of IL-2 solution (8000 U/ml) to a final concentration of 20 U/ml. The blank control wells were divided into five wells, and 100 μl of the medium was added, and 0.5 μl of IL-2 solution (8000 U/ml) was added or not. In addition, zero wells and five wells were added, and 200 μl of medium was added. 37°C, 5% CO2 incubator for 3 days, add WST-1 20ul/well 4 hours before the end of culture, continue culture for 4h, stop the culture, shake the 96-well plate on a shaker for one minute, to fully mix The test system is conditioned. The absorbance (A) value of each well was measured at an ED 450 nm enzyme-linked immunosorbent detector. Based on the absorbance (A) values, the PBMC proliferation rate was calculated as the stimulation index (SI), SI = stimulating the cell hole A450 value / without stimulating the cell hole A450 value.

## Results and Discussion

The hGM-CSF gene HepG2 vaccine and the wild-type HepG2 vaccine were successfully prepared. RT-PCR results can amplify a band of about 410 bp, which is an internal reference. Both transgenic HepG2 cells irradiated with 60Co and unirradiated transgenic HepG2 cells amplified a specific band of about 260 bp, indicating that hGMCSF mRNA was transcribed. There was no specific 260 bp band in wild-type HepG2 cells intermittently irradiated with 60Co and without 60Co intermittent irradiation (Fig. 1). The results of ELISA showed that HepG2 cells transfected with hCo-SF gene by 60Co intermittently secreted hGMCSF, and the secretion amount was 197.28±38.53ng/10E6cells every 24 hours.

**Figure 1.**
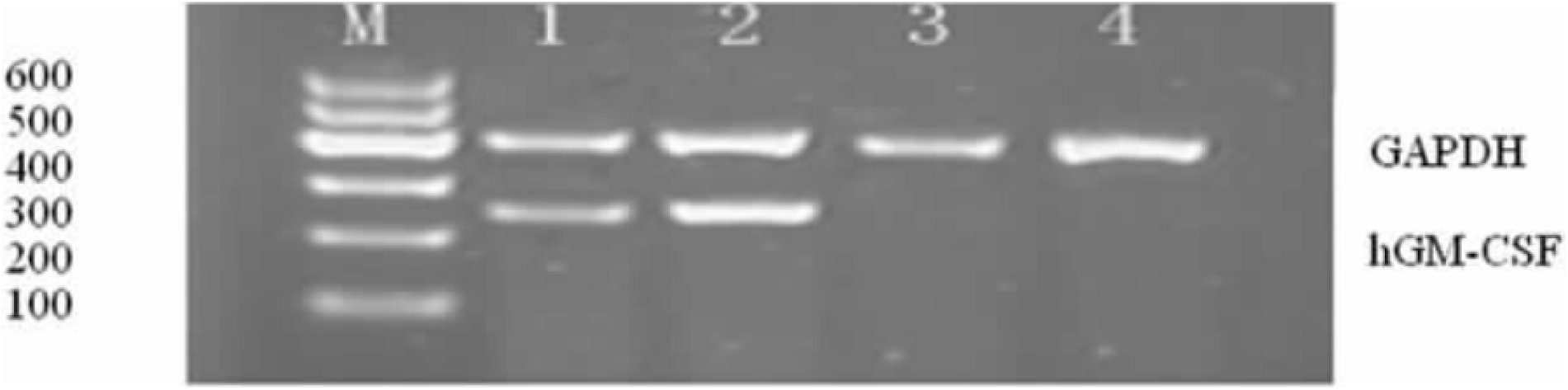
Gel electrophoresis analysis

At the end of the 3-day culture period, the A values of the two blank control PBMCs were 0.398±0.008 and 0.375±0.010, respectively, and the difference was not statistically significant (P>0.05), indicating that 20 U/mL of IL-2 could not be separated. Induction of PBMC proliferation. When the ratio of induced cells (Inducer, I) and effector cells (Effector, E) (I: E) were 1:40, 1:10, 1:20, 1:5, respectively, IL-2 (20 U/) was added. Under the conditions of mL), the results of transgenic HepG2 vaccine and wild-type HepG2 vaccine-stimulated cells to induce PBMC proliferation were as follows (Table 1) (Fig. 2).

**Figure 2.**
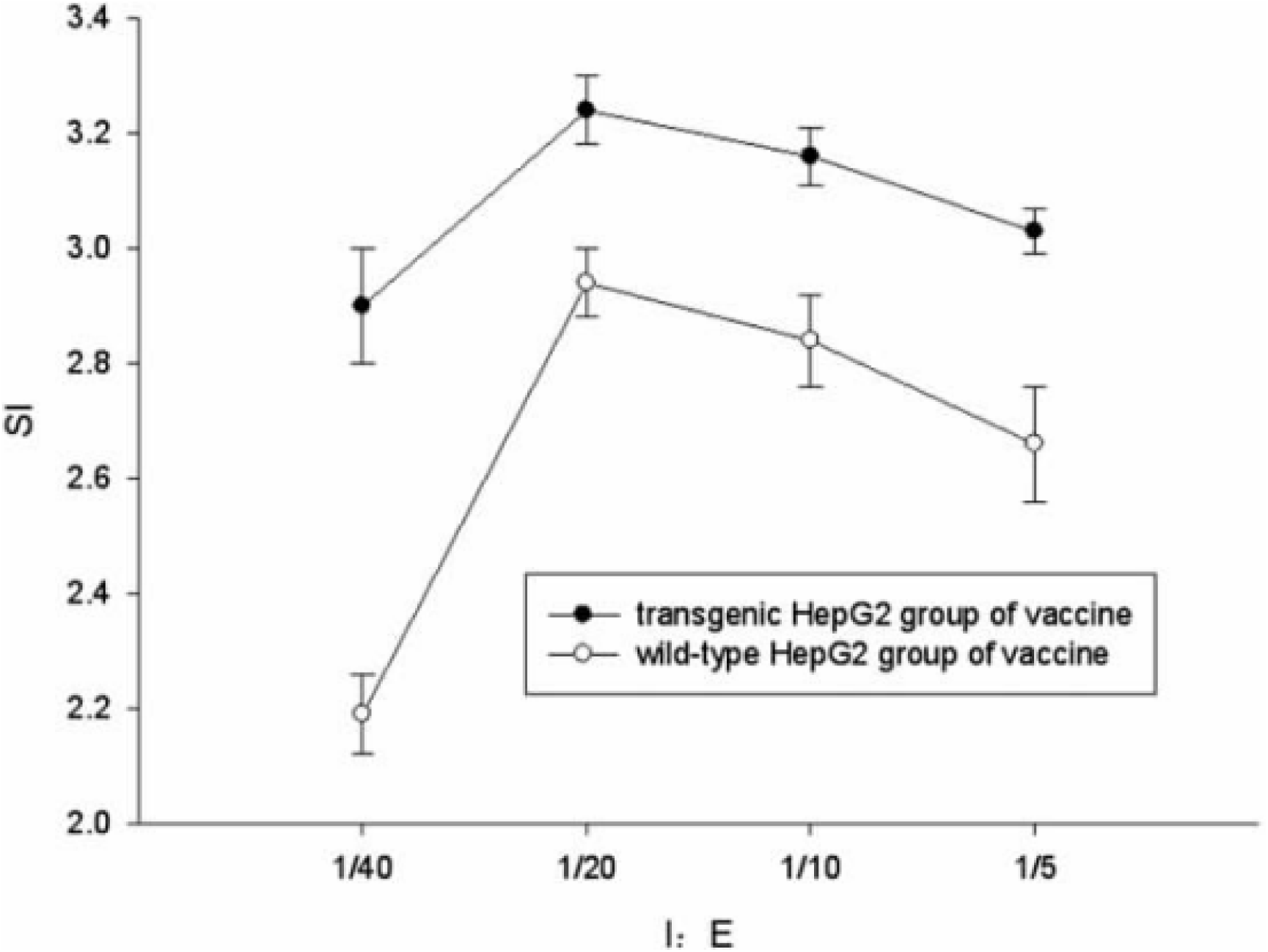
Release kinetics

From the above, we can see that the proliferation of PBMC cells in the transgenic HepG2 vaccine group was higher than that in the wild-type vaccine group A, and the proliferation rate was statistically significant (P<0.05). It indicates that the HepG2 vaccine has enhanced immunogenicity after transduction into the hGMCSF gene, and its ability to induce PBMC proliferation is enhanced. When the I:E value is 1:20, the ability of the transgenic HepG2 vaccine and the wild-type HepG2 vaccine to induce PBMC proliferation has the greatest effect, so the concentration ratio of PBMC induced by the following groups of vaccines was selected.

At the end of the 3-day culture period, the percentage of CD4+ cells in the blank control group A and the blank control group B were (32.07±2.53)%, (30.44±2.02)%, and the percentage of CD8+ cells were respectively (by flow cytometry analysis). 23.25± 2.11)%, (23.03±2.48)%, both P>0.05, indicating that 20 U/mL IL-2 could not induce the percentage change of CD4+ cells and CD8+ cells in PBMC alone. The percentages of CD4+ cells in PBMC of transgenic vaccine group, wild type vaccine group A and wild type vaccine group B were (58.72±2.55)%, (43.43±2.29)%, (50.07±3.20)%, respectively, and the percentage of CD8+ cells were (35.14±2.05)%, (29.41±1.73)%, (32.17±1.58)%, the percentages were higher than the corresponding cells in the blank control group A, all P<0.05. Moreover, the ratio of CD4+ cell positive ratio and CD8+ cell positive ratio in PBMC of transgenic vaccine group was higher than that of wild type vaccine group A and wild type vaccine group B, P<0.05 (Table 2) (Fig. 3).

**Figure 3.**
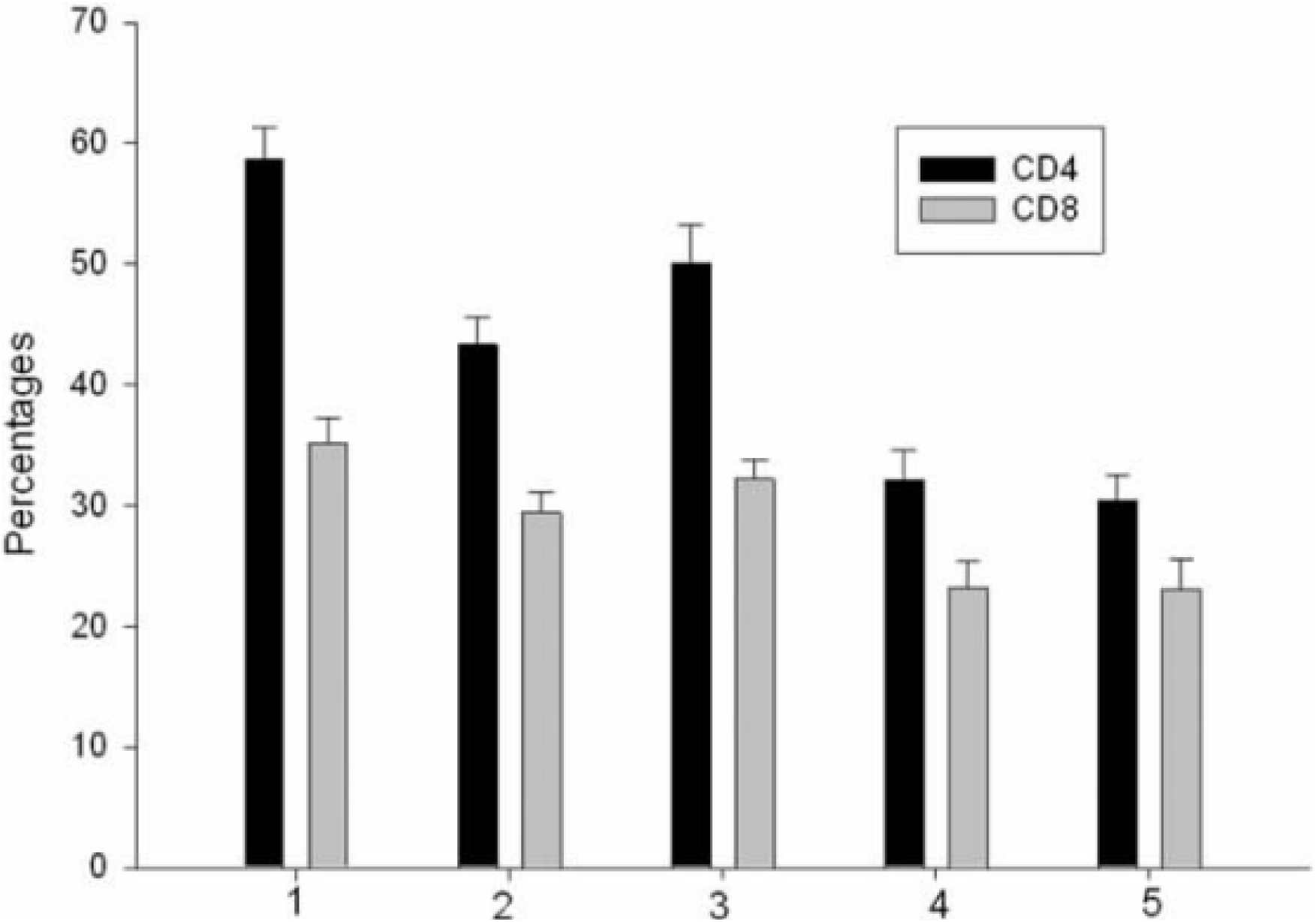
Toxicity assessment

At the end of the 3-day culture period, the levels of IFN-γ in the supernatant of PBMC cultures of the two blank control groups were 757.25 ± 131.91 pg/ml and 692.87 ± 129.24 pg/ml, P>0.05, and the difference was not statistically significant. It was shown that 20 U/mL of IL-2 could not induce PBMC to secrete IFN-γ alone. The contents of IFN-γ in the supernatant of PBMC in transgenic vaccine group, wild type vaccine group A and wild type vaccine group B were 1947.10 ± 133.69 pg/ml, 1032.66 ± 104.79 pg/ml, and 1334.41 ± 159.03 pg/ml, respectively. The IFN-γ content in the supernatant of the vaccine group PBMC was higher than that of the blank control group A, the wild type vaccine group A and the wild type vaccine group B, both P < 0.05 (Table 3) (Fig. 4).

**Figure 4.**
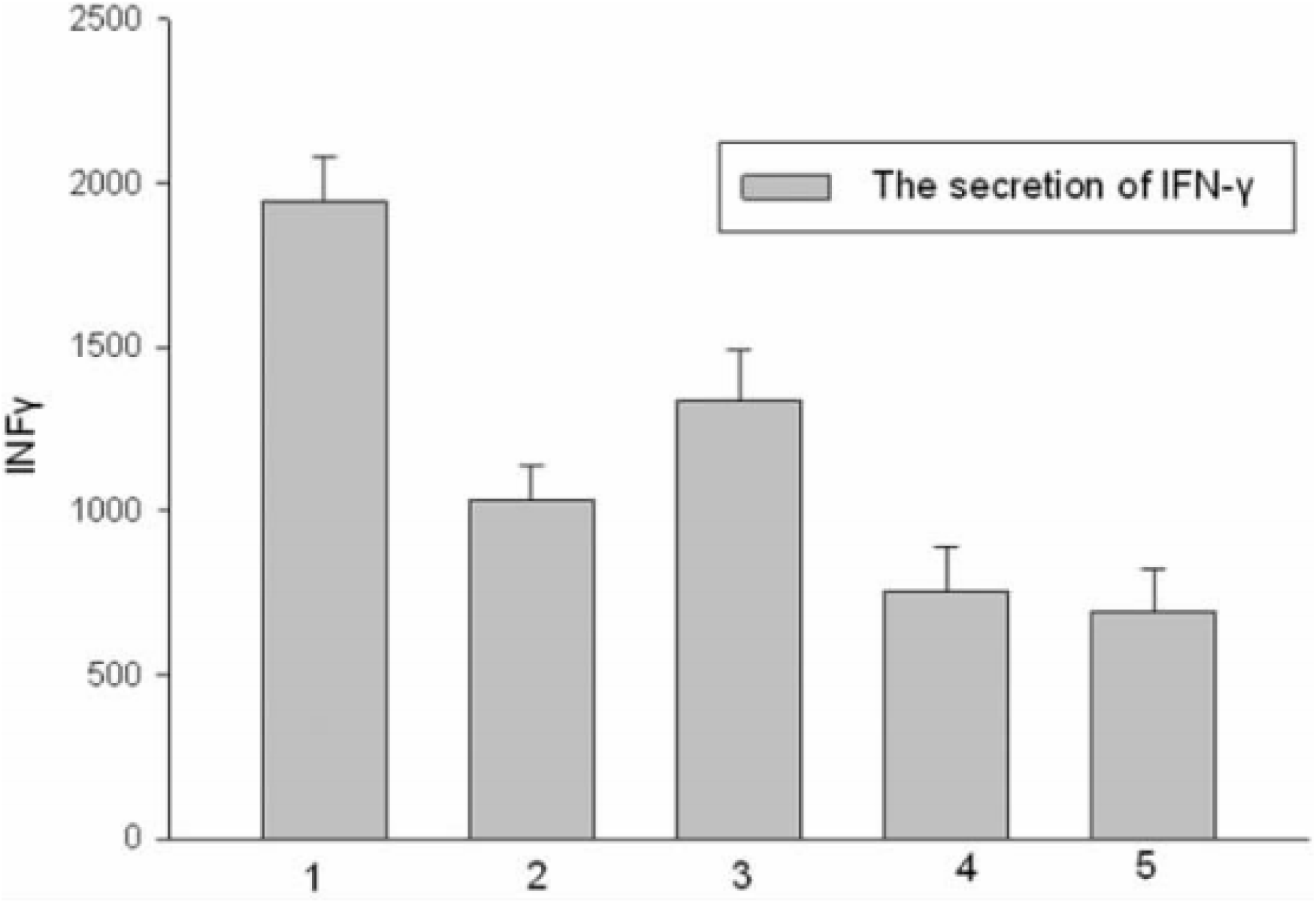
Cellular responses analyzed by ELISA

At the end of the 5-day culture period, the killing rates of PBMCs in the transgenic vaccine group to HepG2 cells were 56.26±4.56, 62.74±, respectively, when the ratio of target to target (E:T) was 12.5: 1,25:1 and 50:1, respectively. 4.82,79.29±4.68, its killing ability was positively correlated with the number of effector cells, higher than the killing rate of blank control group, wild type vaccine group A and wild type vaccine group B, all P<0.05, the difference was statistically significant. The killing rate of HepG2 cells in wild-type vaccine group B was higher than that in wild-type vaccine group A, but P>0.05, the difference was not statistically significant (Table 4) (Fig. 5).

Tumor vaccine is a new type of tumor treatment method that uses tumor cells, tumor cell lysates or tumor antigens to stimulate the body’s immune system to produce specific anti-tumor immune effects. However, autologous and allogeneic whole-cell vaccines stimulate the immune response to a very limited extent, and improve tumors. The exemptive preparation of highly efficient tumor vaccines is the key to immunotherapy. Tumor genetic engineering vaccines use genetic recombination technology to introduce different target genes such as cytokines, helper stimulating molecules, and MHC class I antigen molecules into cells [7], thereby changing the local immune environment and enhancing antigen-presenting cells and tumor specificity. T cell activity. GM-CSF is one of the 33 most potent products evaluated so far [8]. GM-CSF is a cytokine with multiple immunoregulatory functions. Cells modified by GM-CSF gene can express GMCSF for a long time, which can attract a large number of antigen presenting cells to the injection site. These cells capture “tumor cells” here. The tumor antigen, which differentiates, activates, and matures under the action of GM-CSF, presents the decomposed antigen to T lymphocytes, and finally produces activated T lymphocytes capable of killing tumor cells [9]. GM-CSF gene-modified tumor vaccines have shown potential clinical value in experimental and clinical studies of melanoma [10], prostate cancer [11], lung cancer [12], and renal cancer [13]^1-14^. However, GM-CSF gene-modified liver cancer vaccine has not been reported^15^.

On the basis of previous experiments, we successfully prepared transgenic HepG2 cell vaccine by transfecting hGM-CSF gene with HA nanoparticle carrier, and induced PBMC in human peripheral blood mononuclear cells in vitro to observe the proliferation, differentiation and HepG2 cells of PBMC after induction. The killing effect. From the experimental results, the transgenic HepG2 cell vaccine can promote the proliferation of PBMC, up-regulate the expression of CD4+ and CD8+ cells, increase the secretion of INF-γ, and increase its killing effect on HepG2 cells. The immune effect is not only stronger than that of wild-type HepG2^16-20^. The vaccine group and the blank control group were superior to the wild-type vaccine and the equivalent dose of the hGM-CSF mixed group, suggesting that the HepG2 cell vaccine transfected with the hGM-CSF gene induces an increase in antitumor effect in addition to the local effect of GM-CSF secretion. It is also possible that changes in the structure of the antigen after transfer into the hGM-CSF gene are associated with an increase in its immunogenicity. We also noticed the presence of immunodeficiency in patients with liver cancer [14], but due to the difficulty in collecting blood samples, this experiment failed to use peripheral blood in patients after liver cancer surgery; at the same time, the in vitro environment and the in vivo environment are also different. Therefore, further clinical and experimental observations of the vaccine are required after assessing its safety. In conclusion, the HA nanocarrier-mediated HepG2 cell vaccine transfected with the hGM-CSF gene showed a good immune effect in vitro and is expected to be a new approach for the treatment and prevention of HCC metastasis.

## Conclusion

HA nanoparticle-mediated transfection of hGM-CSF gene can increase the immunogenicity of HepG2 cell vaccine, transfecting hGM-CSF gene HepG2 cells The vaccine can effectively induce PBMC proliferation, differentiation, increase the secretion of INF-γ, and enhance its killing effect on HepG2 cells.

